# *alms1* mutant zebrafish do not show hair cell phenotypes seen in other cilia mutants

**DOI:** 10.1101/2020.11.13.381954

**Authors:** Lauren Parkinson, Tamara M. Stawicki

## Abstract

Multiple cilia-associated genes have been shown to affect hair cells in zebrafish (*Danio rerio),* including the human deafness gene *dcdc2,* the radial spoke gene *rsph9,* and multiple intraflagellar transport (IFT) and transition zone genes. Recently a zebrafish *alms1* mutant was generated. The *ALMS1* gene is the gene mutated in the ciliopathy Alström Syndrome a disease that causes hearing loss among other symptoms. The hearing loss seen in Alström Syndrome may be due in part to hair cell defects as *Alms1* mutant mice show stereocilia polarity defects and a loss of hair cells. Hair cell loss is also seen in postmortem analysis of Alström patients. The zebrafish *alms1* mutant has metabolic defects similar to those seen in Alström syndrome and *Alms1* mutant mice. We wished to investigate if it also had hair cell defects. We, however, failed to find any hair cell related phenotypes in *alms1* mutant zebrafish. They had normal lateral line hair cell numbers as both larvae and adults and normal kinocilia formation. They also showed grossly normal swimming behavior, response to vibrational stimuli, and FM1-43 loading. Mutants also showed a normal degree of sensitivity to both short-term neomycin and long-term gentamicin treatment. These results indicate that cilia-associated genes differentially affect different hair cell types.

## INTRODUCTION

Hearing and balance disorders are common sensory disorders (Agrawal *et al.* 2009; Goman and Lin 2016). There is a significant genetic component to these disorders, with over 50% of congenital hearing loss seen in newborns being hereditary (Delmaghani and El-Amraoui 2020). To date, 121 nonsyndromic hearing loss genes have been identified in addition to many syndromic hearing loss genes (Van Camp and Smith). While most of these genes are only implicated in hearing loss, mutations in a small subset also cause balance disorders (Eppsteiner and Smith 2011; Gallego-Martinez *et al.* 2018). Genetic mutations causing hearing loss can affect various processes important for hearing. A number of these mutations specifically affect the development and function of the sensory hair cells responsible for hearing (Delmaghani and El-Amraoui 2020).

Zebrafish have evolved as a useful model to study the genetics of hair cell function. Zebrafish are powerful genetic models and have hair cells on the surface of their bodies as part of the lateral line system, which allows for the observation of hair cells in an intact *in vivo* model. Researchers have carried out large-scale genetic screens to identify zebrafish mutants with auditory or vestibular defects (Malicki *et al.* 1996; Whitfield *et al.* 1996; Nicolson *et al.* 1998) as well as generated zebrafish models of known deafness genes (Blanco-Sánchez *et al.* 2017; Vona *et al.* 2020).

One class of genes that have been shown to influence hair cells in humans and zebrafish are cilia-associated genes. Multiple ciliopathies, genetic disorders with mutations in cilia-associated genes, have been associated with hearing loss, including Usher Syndrome, Alström Syndrome, and Bardet-Biedl Syndrome (BBS) (Alstrom *et al.* 1959; Ross *et al.* 2005; Reiners *et al.* 2006; Girard and Petrovsky 2011; Esposito *et al.* 2017). A number of Usher Syndrome genes have also been shown to affect hair cell function in zebrafish (Ernest *et al.* 2000; Söllner *et al.* 2004; Seiler *et al.* 2005; Ebermann *et al.* 2010; Riazuddin *et al.* 2012; Ogun and Zallocchi 2014). Other cilia genes associated with hearing loss in humans include *DCDC2, CDC14A, CCDC114,* and the basal body genes *CEP78* and *CEP250* (Khateb *et al.* 2014; Grati *et al.* 2015; Namburi *et al.* 2016; Delmaghani *et al.* 2016; Nikopoulos *et al.* 2016; Li *et al.* 2018). While some of these genes, such as *dcdc2,* are similarly necessary for hair cell function in zebrafish (Grati *et al.* 2015), others such as *cdc14A* do not cause zebrafish hair cell phenotypes when mutated (Imtiaz *et al.* 2018). There have also been a number of cilia-associated genes that have not been implicated in human deafness but have been shown to cause hair cell phenotypes when disrupted in zebrafish. Mutations in intraflagellar transport (IFT) genes, genes necessary for the transport of proteins along cilia, in zebrafish have been shown to decrease hair cell numbers, lead to resistance to aminoglycoside-induced hair cell death, and to cause defects in FM1-43 uptake and early reverse polarity hair cell mechanotransduction activity (Tsujikawa and Malicki 2004; Kindt *et al.* 2012; Stawicki *et al.* 2016, 2018). Mutations in transition zone genes, genes that act as gatekeepers for proteins exiting and entering cilia, have also been shown to cause resistance to aminoglycoside-induced hair cell death but to have normal FM1-43 uptake and control hair cell numbers (Owens *et al.* 2008; Stawicki *et al.* 2016), and a mutation in *rsph9,* a radial spoke gene in motile cilia, caused a reduced initiation of the startle response to acoustic stimuli in zebrafish suggesting a potential hair cell defect (Sedykh *et al.* 2016).

Recently a zebrafish *alms1* mutant line was generated (Nesmith *et al.* 2019). *ALMS1* is the gene responsible for Alström Syndrome, a ciliopathy characterized by hearing and vision loss in addition to obesity and diabetes (Alstrom *et al.* 1959; Hearn *et al.* 2002; Collin *et al.* 2002). Dilated cardiomyopathy, hypertriglyceridemia, gastrointestinal disturbances, neurological symptoms, and liver and kidney dysfunction have also been observed in some patients (Marshall *et al.* 2005). ALMS1 localizes to the basal body at the base of cilia leading to the classification of Alström Syndrome as a ciliopathy (Hearn *et al.* 2005). Hearing loss in Alström Syndrome patients usually begins in childhood and is progressive (Marshall *et al.* 1997, 2005; Lindsey *et al.* 2017) with patients also showing abnormal distortion product otoacoustic emissions (DPOAEs) (Bahmad *et al.* 2014; Lindsey *et al.* 2017) Vestibular defects have not been observed. Postmortem analysis of patient auditory tissue shows degeneration of the organ of Corti, including a loss of hair cells. Additionally, there is atrophy of the stria vascularis and spiral ligament and degeneration of the spiral ganglion. In contrast to the auditory tissue, vestibular epithelium looked normal in these patients, in agreement with the lack of vestibular symptoms (Nadol *et al.* 2015). Mouse *Alms1* mutants show similar phenotypes as human patients, including obesity, retinal dysfunction, hyperinsulinemia, and delayed onset hearing loss (Collin *et al.* 2005; Arsov *et al.* 2006; Jagger *et al.* 2011). Hair cells in *Alms1* mutant mice show misshapen stereocilia bundles and polarity defects and a loss of outer hair cells with age. Strial atrophy is also seen similar to what was observed in human postmortem samples (Jagger *et al.* 2011; Nadol *et al.* 2015). Similar to mouse mutants, zebrafish *alms1* mutants share many phenotypes with Alström Syndrome patients, including retinal degeneration, kidney and cardiac defects, increased fat disposition in the liver, a propensity for obesity, hyperinsulinemia, and glucose response defects (Nesmith *et al.* 2019). However, hair cells and the auditory and vestibular systems of these fish have not been examined.

We wished to examine *alms1* mutant zebrafish to see if they showed hair cell phenotypes similar to mammalian *Alms1* mutants and other zebrafish cilia-associated gene mutants. We found that *alms1* mutant zebrafish had normal cilia formation in lateral line hair cells and other ciliated cells similar to what has been observed in mammalian *Alms1* mutants. We also failed to find any hair cell phenotypes in these mutants. They showed grossly normal audiovestibular behavior and sensitivity to aminoglycosides as both larvae and adults. They also showed normal FM1-43 uptake into hair cells. These results and the lack of vestibular defects seen in mammals following *Alms1* mutations suggest a specific role for *Alms1* in the auditory system rather than more global hair cell function. It also shows that cilia-associated genes have distinct roles in different hair cell types.

## MATERIALS AND METHODS

### Animals

Experiments used either 5 days post-fertilization (dpf) *Danio rerio* (zebrafish) larvae or adult zebrafish over three months of age. We used the previously described *alms1*^*umd2*^ mutant line, which causes a premature stop codon (Nesmith *et al.* 2019). Mutant larvae were generated by either crossing two heterozygous animals together, or one heterozygous animal to a homozygous mutant, and comparing homozygous mutants to either wild-type or heterozygous siblings born at the same time. For the experiment on adult zebrafish, homozygous mutants were incrossed to generate the mutant animals, and homozygous wild-type siblings of those mutants were crossed to heterozygous wild-type siblings to generate the wild-type controls. Larvae were raised in embryo media (EM) consisting of 1 mM MgSO_4_, 150 μM KH_2_PO_4_, 42 μM Na_2_HPO_4_, 1 mM CaCl_2_, 500 μM KCl, 15 mM NaCl, and 714 μM NaHCO_3_ and housed in an incubator maintained at 28.5°C with a 14/10 hour light/dark cycle. The Lafayette College Institution Animal Care and Use Committee approved all experiments.

### Antibody labeling

Zebrafish larvae and fins used for antibody labeling were fixed for two hours at room temperature in 4% paraformaldehyde. Antibody labeling was carried out as previously described (Stawicki *et al.* 2014). Cilia were labeled with a mouse anti-acetylated tubulin primary antibody (Millipore-Sigma, T7451) diluted at 1:1,000 in antibody block. Larvae used for hair cell counts were labeled with a rabbit anti-parvalbumin primary antibody (ThermoFisher, PA1-933) diluted at 1:1,000 in antibody block. Fins used for hair cell counts were labeled with both the parvalbumin antibody and a mouse anti-otoferlin antibody (Developmental Studies Hybridoma Bank, HCS-1) diluted at 1:00 in antibody block.

### FM1-43 Uptake

Fish were treated with 2.25 μM FM 1-43FX (ThermoFisher, F35355) for 1 minute in EM, washed 3 times in EM, and then anesthetized with MS-222/Tricaine-S (Pentair Aquatic Eco-Systems, TRS1) for imaging. Fish were imaged on a Zeiss LSM800 confocal microscope. For each fish, a single neuromast was imaged by taking a stack of 5 1 μm optical sections. Image analysis was carried out in Fiji. A maximum projection image was made of each stack of images, and the average fluorescent intensity of the cell bodies of the entire neuromast was measured. Additionally, the average fluorescent intensity of an area in the background of the image was measured. Finally, the value for the fluorescence of the neuromast was divided by the value of the background fluorescence. Larvae were euthanized and genotyped after imaging.

### Aminoglycoside Treatment and Hair Cell Counts

Fish were treated with either neomycin solution (Sigma-Aldrich, N1142) or gentamicin solution (Sigma-Aldrich G1272) at the indicated doses diluted in EM. For all neomycin treatment experiments, fish were treated with neomycin for 30 minutes, washed 3 times in either plain EM for larvae or plain system water for adult fish, left to recover in the third wash for one hour, and then tissue was collected for antibody labeling. In the case of larvae, whole larvae were fixed, whereas for adult fish, the fish were anesthetized, and their caudal fins were amputated and fixed. For gentamicin treatment, larvae were treated with gentamicin for 6 hours, washed 3 times in plain EM, and then fixed for antibody labeling. For adult zebrafish, 6 neuromasts on each fin were counted using the HCS-1 stain, and an average hair cell/neuromast number was generated for each animal. For larvae, the OP1, M2, IO4, MI2, and MI1 neuromasts (Raible and Kruse 2000) were counted using the parvalbumin stain, and again an average number of hair cells/neuromast number was generated for each animal. Larvae were genotyped after counting was complete. All hair cell counts were carried out on an Accu-Scope EXC-350 microscope.

### Statistical analysis

All statistics were calculated using GraphPad Prism software (version 6.0).

## RESULTS

### Cilia morphology is normal in *alms1* mutants

There have previously been conflicting results regarding whether ALMS1 plays a role in cilia formation and maintenance. Cilia in *Alms1* mutant mice and fibroblasts isolated from Alström syndrome patients appear grossly normal (Collin *et al.* 2005, 2012; Hearn *et al.* 2005; Li *et al.* 2007; Chen *et al.* 2017) even when there is no visible antibody labeling for ALMS1 protein (Chen *et al.* 2017). However, RNAi knockdown of *Alms1* in cultured cells results in abnormal and stunted cilia (Li *et al.* 2007; Graser *et al.* 2007). To test what cilia morphology looked like in zebrafish *alms1* mutants, we stained 5dpf zebrafish larvae with acetylated tubulin. We found that cilia morphology looked grossly normal in both hair cells of the lateral line and cells of the olfactory pit in *alms1* mutants (Figure 1).

**Figure 1:**
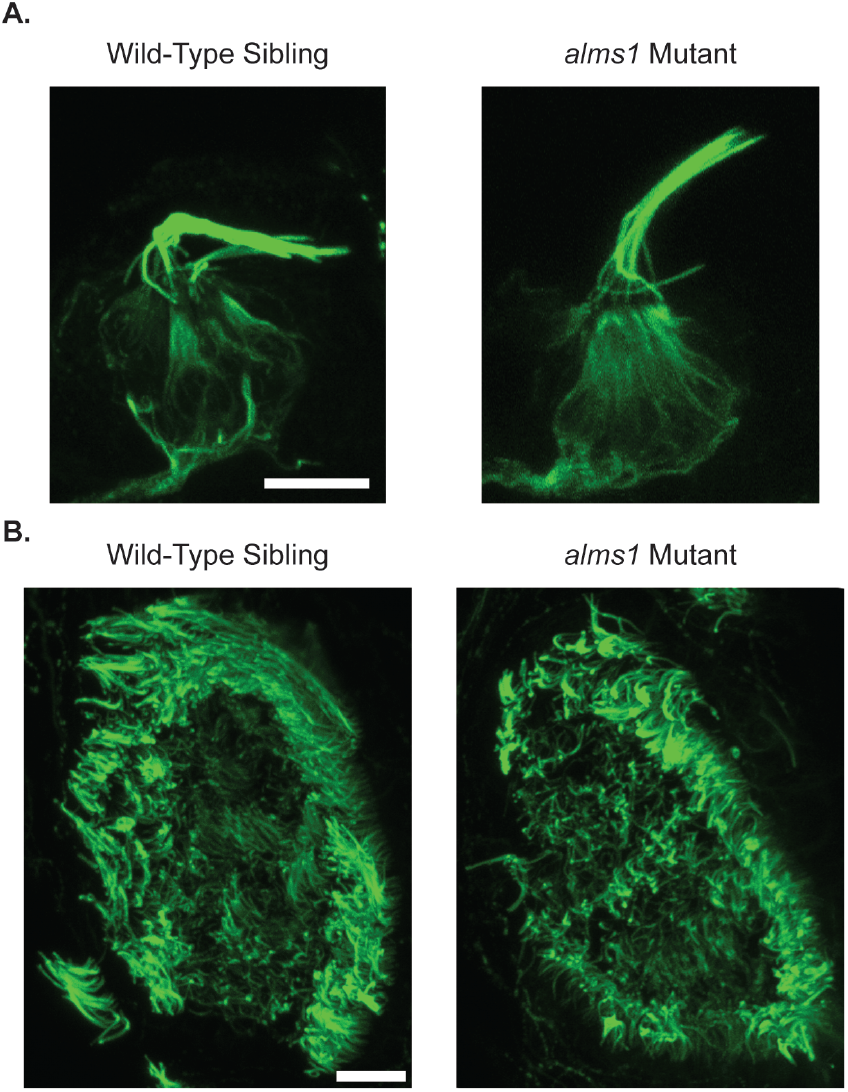
Cilia formation is normal in *alms1* mutants. (A) Representative images of the OC1 neuromast stained with acetylated tubulin in wild-type (left) and *alms1* mutant (right) 5dpf zebrafish larvae. Kinocilia appear grossly normal in *alms1* mutant fish. (B) Representative images of the olfactory pit stained with acetylated tubulin in wild-type (left) and *alms1* mutant (right) 5dpf zebrafish larvae. Again cilia appear grossly normal in *alms1* mutants. Scale bar = 10 μm.

### *alms1* mutants do not show mechanotransduction defect phenotypes

Zebrafish mechanotransduction mutants show several characteristic behavioral phenotypes, including a failure to remain upright, circling behavior when swimming, and a failure to respond to acoustic-vibrational stimuli (Nicolson *et al.* 1998). It has previously been shown that zebrafish morphants of another cilia-associated deafness gene, *dcdc2,* show similar behavioral defects (Grati *et al.* 2015). We, however, failed to observe any of these phenotypes in *alms1* mutant zebrafish. *alms1* 5dpf mutant larvae remain upright, and those without body morphology defects showed a grossly normal escape response following a vibrational stimulus generated by taping on their dish (Movie S1). Adult homozygous mutants also showed normal swimming behavior (Movie S2). The rapid uptake of FM1-43 is also reduced or eliminated when hair cell mechanotransduction activity is impaired (Seiler and Nicolson 1999; Gale *et al.* 2001; Meyers *et al.* 2003). It has previously been shown that other cilia gene mutants, specifically IFT mutants, show reduced levels of FM1-43 loading into lateral line hair cells (Stawicki *et al.* 2016, 2018). However, we failed to observe any significant differences in FM1-43 loading in *alms1* mutants compared to wild-type siblings (Figure 2).

**Figure 2:**
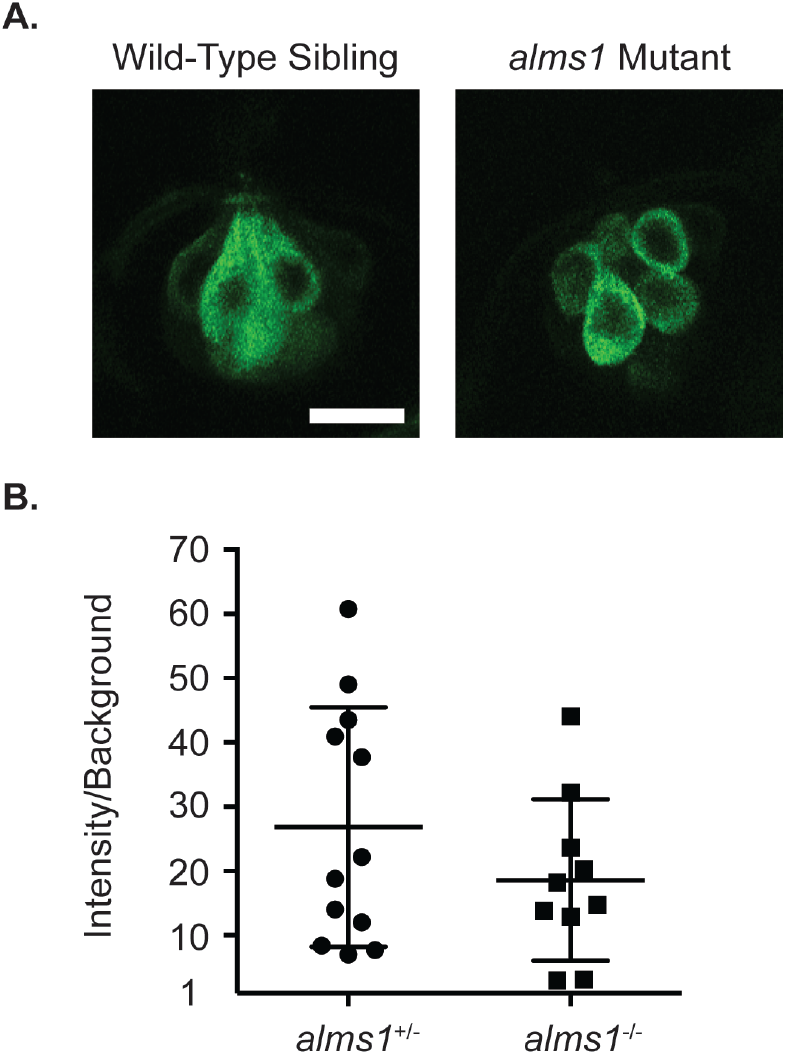
FM1-43 loading is normal in *alms1* mutants. (A) Representative images of neuromasts from wild-type siblings (left) and *alms1* mutants (right) treated with FM1-43. Scale bar = 10 μm. (B) Quantification of the fluorescent intensity of FM1-43 in neuromasts of heterozygous wild-type siblings and homozygous *alms1* mutants. There was no significant difference between the two groups by an unpaired t-test (p = 0.2474). Individual data points are shown along with the mean and standard deviation of the data.

### *alms1* mutants are not resistant to aminoglycoside-induced hair cell death

It has previously been shown that some zebrafish cilia-associated gene mutants are resistant to aminoglycoside-induced hair cell death, specifically transition zone, including the basal body gene *cep290,* and IFT gene mutants (Owens *et al.* 2008; Stawicki *et al.* 2016, 2018). To test if this was also the case for *alms1* mutants, we used both short-term neomycin and long-term gentamicin treatment paradigms and quantified hair cell number in 5dpf zebrafish larvae. We failed to see any significant differences between wild-type mutants and homozygous mutants in either control conditions or following aminoglycoside treatment (Figure 3A, C). Many cilia-associated genes are maternally expressed in zebrafish, meaning a heterozygous mother loads wild-type RNA into the egg, and this can mask mutant phenotypes early in development (Huang and Schier 2009; Duldulao *et al.* 2009; Cao *et al.* 2010). Therefore, we wished to test if this was happening in *alms1* mutants by generating mutant larvae out of both heterozygous and homozygous mutant mothers and treating them with neomycin. We again failed to find any significant differences between the different genotypes (Figure 3B).

**Figure 3:**
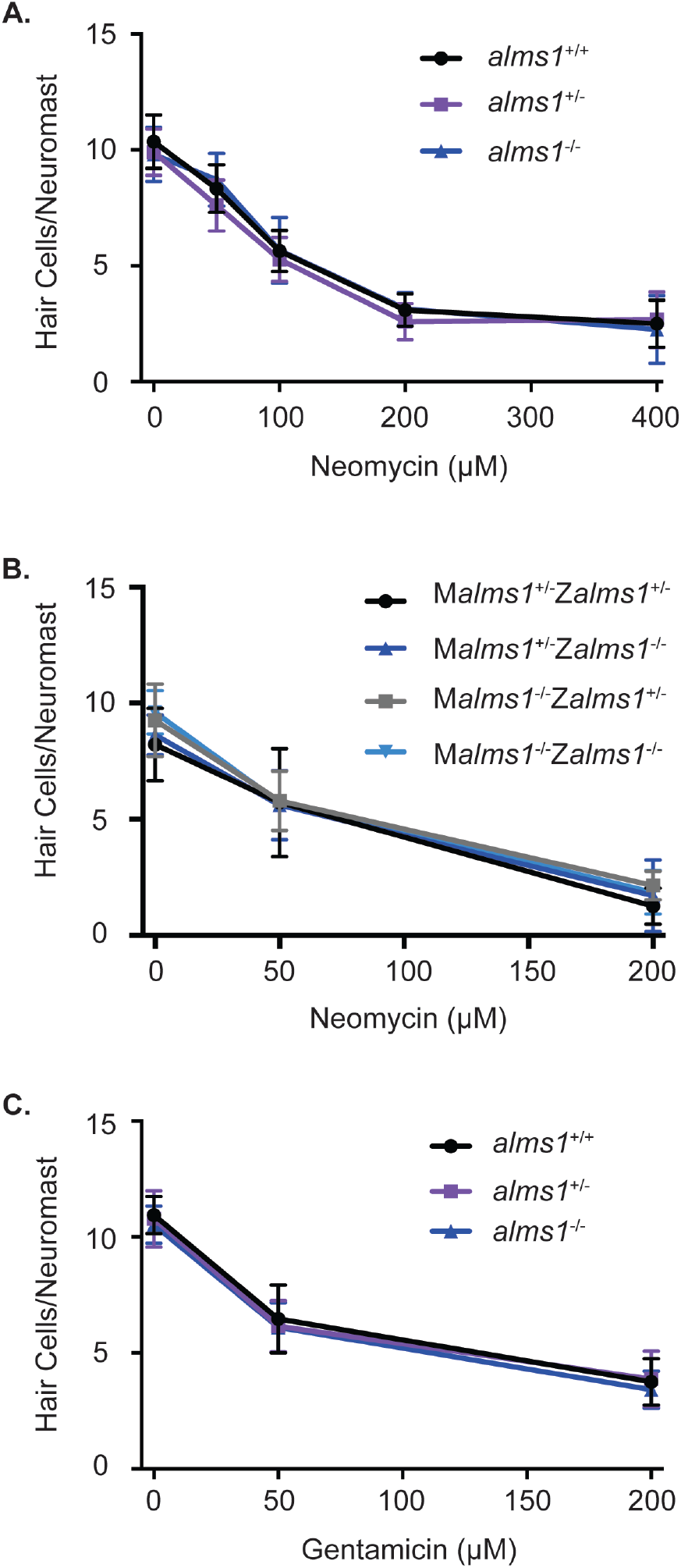
*alms1* mutant larvae are not resistant to aminoglycoside-induced hair cell death. (A) Dose-response curve for wild-type siblings and *alms1* mutants from an incross of heterozygous parents treated with 0-400 μM neomycin for 30 minutes with a one-hour recovery time. n = 7-19 fish per neomycin dose for homozygous wild-type and mutants and 20-25 for heterozygous fish. There were no significant differences due to genotype (p= 0.1273) or the interaction between genotype and neomycin (p = 0.3939) by two-way ANOVA. (B) Dose-response curve for fish that were either heterozygous or homozygous for the *alms1* mutation from mothers also either heterozygous or homozygous for the *alms1* mutation treated with 0-200 μM neomycin for 30 minutes with a one-hour recovery time. M = maternal genotype and Z = zygotic genotype. n = 7-16 fish per neomycin dose for each genotype. There were no significant differences due to genotype (p= 0.8060) or the interaction between genotype and neomycin (p = 0.2024) by two-way ANOVA. (C) Dose-response curve for wild-type siblings and *alms1* mutants from parents heterozygous for the *alms1* mutation treated with 0-200 μM gentamicin for 6 hours. n = 10-14 fish per gentamicin dose for homozygous wild-type and mutants and 21-25 for heterozygous fish. There were no significant differences due to genotype (p= 0.9401) or the interaction between genotype and neomycin (p = 0.2850) by two-way ANOVA. Data are shown as mean +/− standard deviation.

In human Alström syndrome, patients’ hearing loss is usually not seen until later in childhood (Michaud *et al.* 1996; Marshall *et al.* 2005; Lindsey *et al.* 2017). Likewise, hearing defects in *Alms1* mutant mice show a delayed onset (Collin *et al.* 2005; Arsov *et al.* 2006; Jagger *et al.* 2011). Therefore, we wished to see if adult zebrafish were resistant to aminoglycoside-induced hair cell death even though larvae were not. We treated fish slightly over 3 months of age that appeared to be mature adults (Movie S2) with neomycin. We again failed to find any significant difference between homozygous mutants and related wild-type fish in either control lateral line hair cell numbers or hair cell numbers following neomycin treatment (Figure 4).

**Figure 4:**
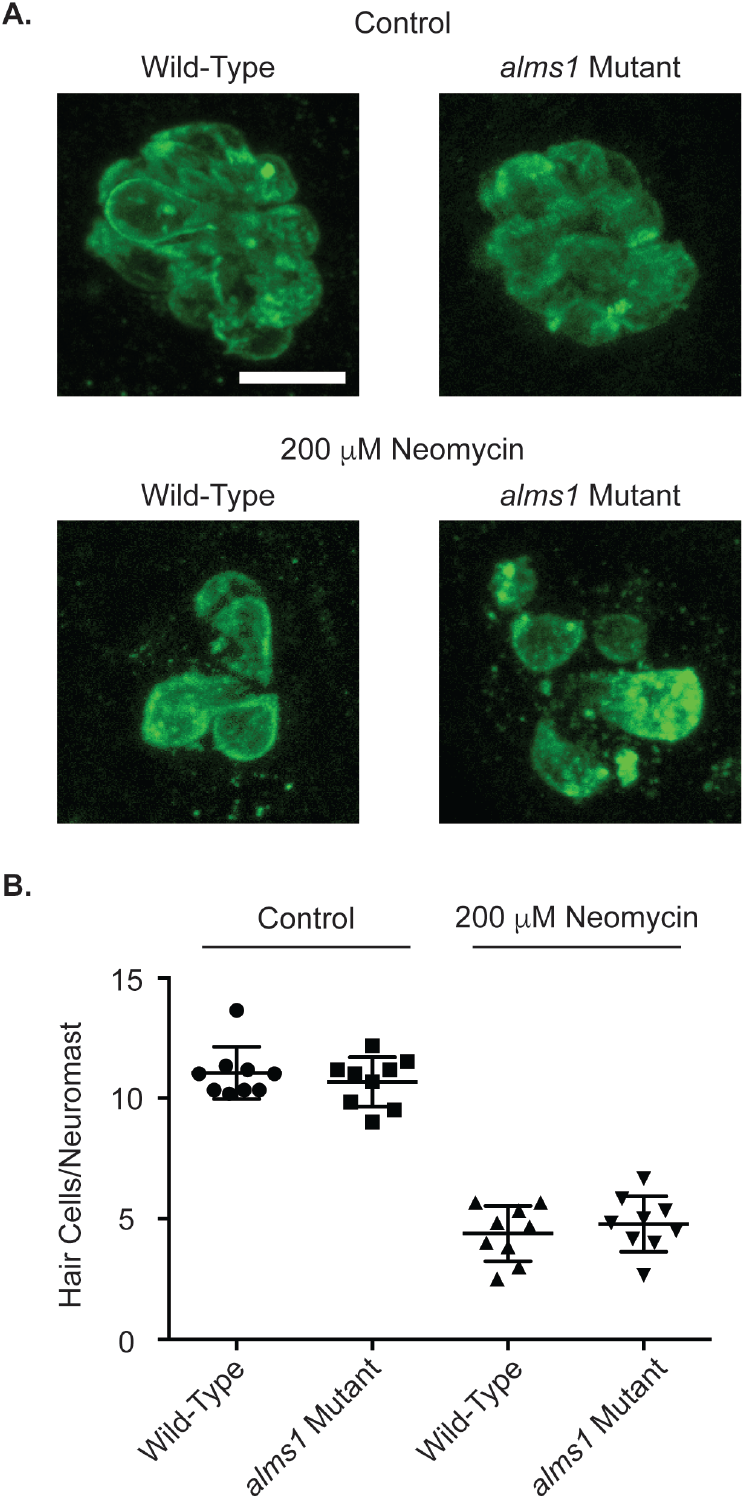
Adult *alms1* mutant fish are not resistant to neomycin-induced hair cell death. (A) Representative images of neuromasts from control (top) and neomycin treated (bottom) fish from wild-type (left) and *alms1* mutant (right) fish. (B) Quantification of hair cells/neuromast either in control conditions or following treatment with 200 μM neomycin. Individual data points are shown along with the mean and standard deviation. There were no significant differences due to genotype (p= 0.3073) or the interaction between genotype and neomycin (p = 0.9784) by two-way ANOVA.

## DISCUSSION

Zebrafish have emerged as a useful model to study hair cell function and death in part due to the external location of hair cells in the lateral line system. Multiple genes associated with both syndromic and nonsyndromic forms of deafness have been shown to cause hair cell related phenotypes in zebrafish (Blanco-Sánchez *et al.* 2017; Vona *et al.* 2020). This is particularly true for genes affecting mechanotransduction or synaptic activity (Nicolson 2015, 2017). However, not all deafness genes have been shown to have phenotypes when mutated in zebrafish, including the cilia-associated gene *cdc114* (Imtiaz *et al.* 2018). We found this also to be the case with *alsm1* where mutants showed grossly normal swimming behavior and response to vibrational stimuli, normal hair cell number in contrast to what has previously been shown in mouse *Alms1* mutants (Jagger *et al.* 2011) and human patients (Nadol *et al.* 2015), and normal FM1-43 uptake and aminoglycoside toxicity of lateral line hair cells in contrast to what has been seen in other zebrafish cilia-associated gene mutants (Owens *et al.* 2008; Stawicki *et al.* 2016, 2018).

There are multiple explanations for the lack of phenotypes we observed. First, it has previously been shown that CRISPR mutants can show genetic compensation that is not present when genes are merely knocked down (Rossi *et al.* 2015). It is also possible that *alms1* might not be playing a role in zebrafish hair cells due to a lack of conservation to the mammalian homolog as much of the N-terminal half of the protein is specific to mammals (Hearn 2019). However, as the zebrafish mutant shows multiple expected phenotypes due to defects in other tissues (Nesmith *et al.* 2019) if any of these issues were responsible for the lack of phenotype we see, it would have to be relatively hair cell specific. We can also not rule out more subtle defects, particularly in inner ear hair cells, that were not observed by the methods we used.

However, we feel the most likely explanation for the lack of observed phenotypes is different roles for *alms1* in different hair cell types. While human Alström patients and *Alms1* mutant mice show hearing defects and defects in auditory hair cells, there have not been any reported vestibular defects. ALMS1 specifically localizes to the centrosome or basal body of cilia (Hearn *et al.* 2005; Jagger *et al.* 2011), and other basal body proteins have likewise been associated with hearing loss but not vestibular dysfunction (Khateb *et al.* 2014; Nikopoulos *et al.* 2016). While different hair cell types have many shared genes, there are also many genes that are auditory or vestibular hair cell specific (Scheffer *et al.* 2015; Burns *et al.* 2015). Comparing lists of highly expressed zebrafish hair cell genes (Erickson and Nicolson 2015; Barta *et al.* 2018) to these lists show overlap with several vestibular hair cell specific genes but no auditory hair cell specific genes. This suggests zebrafish hair cells are more closely related to mammalian vestibular hair cells than auditory hair cells. One of the defects seen in *Alms1* mutant hair cells in mammals, is stereocilia polarity (Jagger *et al.* 2011). This defect has also been shown in auditory hair cells of other cilia-associated gene mutants (Ross *et al.* 2005; Jones *et al.* 2008; May-Simera *et al.* 2009; Sipe and Lu 2011). However, in some cases, these mutants had no polarity defects in vestibular hair cells (Jones *et al.* 2008; Sipe and Lu 2011), and these polarity defects have not been seen in lateral line hair cells of zebrafish cilia-associated gene mutants (Kindt *et al.* 2012; Stawicki *et al.* 2016). Therefore, it appears that in mammalian auditory hair cells where the kinocilia, the true cilia of the cell, is lost early in development (Lim and Anniko 1985) the primary role of cilia-associated genes has evolved to be determining stereocilia polarity early in development. Whereas, in mammalian vestibular hair cells and zebrafish hair cells, where the kinocilia are maintained, these genes may play other functions that are not yet fully elucidated.

## ACKNOWLEDGEMENTS

We thank Jessica Nesmith and Norann Zaghloul for the *alms1* mutant strain, Ivan Cruz for advice on labeling hair cells in adult zebrafish, and Amy Badillo for assistance with zebrafish care.

## Notes

### Competing Interest Statement

The authors have declared no competing interest.

## LITERATURE CITED

Agrawal Y., J. P. Carey, C. C. Della Santina, M. C. Schubert, and L. B. Minor, 2009 Disorders of balance and vestibular function in US adults: Data from the National Health and Nutrition Examination Survey, 2001-2004. Arch. Intern. Med. 169: 938–944. https://doi.org/10.1001/archinternmed.2009.66

Alstrom C., B. Hallgren, L. Nilsson, and H. Asander, 1959 Retinal degeneration combined with obesity, diabetes mellitus and neurogenous deafness: a specific syndrome (not hitherto described) distinct from the Laurence-Moon-Bardet-Biedl syndrome: a clinical, endocrinological and genetic examination based on a lar. Acta Psychiatr Neurol Scand Suppl 129: 1–35.

Arsov T., D. G. Silva, M. K. O’Bryan, A. Sainsbury, N. J. Lee, et al., 2006 Fat aussie - A new Alström syndrome mouse showing a critical role for ALMS1 in obesity, diabetes, and spermatogenesis. Mol. Endocrinol. 20: 1610–1622. https://doi.org/10.1210/me.2005-0494

Bahmad F., C. S. A. Costa, M. S. Teixeira, J. de Barros Filho, L. M. Viana, et al., 2014 Familial Alström syndrome: a rare cause of bilateral progressive hearing loss. Braz. J. Otorhinolaryngol. 80: 99–104. https://doi.org/10.5935/1808-8694.20140023

Barta C. L., H. Liu, L. Chen, K. P. Giffen, Y. Li, et al., 2018 RNA-seq transcriptomic analysis of adult zebrafish inner ear hair cells. Sci. Data 5: 180005. https://doi.org/10.1038/sdata.2018.5

Blanco-Sánchez B., A. Clément, J. B. Phillips, and M. Westerfield, 2017 Zebrafish models of human eye and inner ear diseases. Methods Cell Biol. 138: 415–467. https://doi.org/10.1016/bs.mcb.2016.10.006

Burns J. C., M. C. Kelly, M. Hoa, R. J. Morell, and M. W. Kelley, 2015 Single-cell RNA-Seq resolves cellular complexity in sensory organs from the neonatal inner ear. Nat. Commun. 6: 8557. https://doi.org/10.1038/ncomms9557

Cao Y., A. Park, and Z. Sun, 2010 Intraflagellar transport proteins are essential for cilia formation and for planar cell polarity. J. Am. Soc. Nephrol. 21: 1326–33. https://doi.org/10.1681/ASN.2009091001

Chen J. H., T. Geberhiwot, T. G. Barrett, R. Paisey, and R. K. Semple, 2017 Refining genotype–phenotype correlation in Alström syndrome through study of primary human fibroblasts. Mol. Genet. Genomic Med. 5: 390–404. https://doi.org/10.1002/mgg3.296

Collin G. B., J. D. Marshall, A. Ikeda, W. V. So, I. Russell-Eggitt, et al., 2002 Mutations in ALMS1 cause obesity, type 2 diabetes and neurosensory degeneration in Alström syndrome. Nat. Genet. 31: 74–78. https://doi.org/10.1038/ng867

Collin G. B., E. Cyr, R. Bronson, J. D. Marshall, E. J. Gifford, et al., 2005 Alms1-disrupted mice recapitulate human Alströ m syndrome. Hum. Mol. Genet. 14: 2323–2333. https://doi.org/10.1093/hmg/ddi235

Collin G. B., J. D. Marshall, B. L. King, G. Milan, P. Maffei, et al., 2012 The alström syndrome protein, ALMS1, interacts with α-actinin and components of the endosome recycling pathway. PLoS One 7: e37925. https://doi.org/10.1371/journal.pone.0037925

Delmaghani S., A. Aghaie, Y. Bouyacoub, H. El Hachmi, C. Bonnet, et al., 2016 Mutations in CDC14A, Encoding a Protein Phosphatase Involved in Hair Cell Ciliogenesis, Cause Autosomal-Recessive Severe to Profound Deafness. Am. J. Hum. Genet. 98: 1266–1270. https://doi.org/10.1016/j.ajhg.2016.04.015

Delmaghani S., and A. El-Amraoui, 2020 Inner Ear Gene Therapies Take Off: Current Promises and Future Challenges. J. Clin. Med. 9: 2309. https://doi.org/10.3390/jcm9072309

Duldulao N. A., S. Lee, and Z. Sun, 2009 Cilia localization is essential for in vivo functions of the Joubert syndrome protein Arl13b/Scorpion. Development 136: 4033–42. https://doi.org/10.1242/dev.036350

Ebermann I., J. B. Phillips, M. C. Liebau, R. K. Koenekoop, B. Schermer, et al., 2010 PDZD7 is a modifier of retinal disease and a contributor to digenic Usher syndrome. J. Clin. Invest. 120: 1812–1823. https://doi.org/10.1172/JCI39715

Eppsteiner R. W., and R. J. H. Smith, 2011 Genetic disorders of the vestibular system. Curr. Opin. Otolaryngol. Head Neck Surg. 19: 397–402. https://doi.org/10.1097/MOO.0b013e32834a9852

Erickson T., and T. Nicolson, 2015 Identification of sensory hair-cell transcripts by thiouracil-tagging in zebrafish. BMC Genomics 16: 842. https://doi.org/10.1186/s12864-015-2072-5

Ernest S., G. J. Rauch, P. Haffter, R. Geisler, C. Petit, et al., 2000 Mariner is defective in myosin VIIA: A zebrafish model for human hereditary deafness. Hum. Mol. Genet. 9: 2189–2196. https://doi.org/10.1093/hmg/9.14.2189

Esposito G., F. Testa, M. Zacchia, A. A. Crispo, V. Di Iorio, et al., 2017 Genetic characterization of Italian patients with Bardet-Biedl syndrome and correlation to ocular, renal and audio-vestibular phenotype: Identification of eleven novel pathogenic sequence variants. BMC Med. Genet. 18: 10. https://doi.org/10.1186/s12881-017-0372-0

Gale J. E., W. Marcotti, H. J. Kennedy, C. J. Kros, and G. P. Richardson, 2001 FM1-43 dye behaves as a permeant blocker of the hair-cell mechanotransducer channel. J. Neurosci. 21: 7013–7025.

Gallego-Martinez A., J. M. Espinosa-Sanchez, and J. A. Lopez-Escamez, 2018 Genetic contribution to vestibular diseases. J. Neurol. 265 (Suppl: S29–S34.

Girard D., and N. Petrovsky, 2011 Alström syndrome: Insights into the pathogenesis of metabolic disorders. Nat. Rev. Endocrinol. 7: 77–88.

Goman A. M., and F. R. Lin, 2016 Prevalence of Hearing Loss by Severity in the United States. Am. J. Public Health 106: 1820–1822. https://doi.org/10.2105/AJPH.2016.303299

Graser S., Y. D. Stierhof, S. B. Lavoie, O. S. Gassner, S. Lamla, et al., 2007 Cep164, a novel centriole appendage protein required for primary cilium formation. J. Cell Biol. 179: 321–330. https://doi.org/10.1083/jcb.200707181

Grati M., I. Chakchouk, Q. Ma, M. Bensaid, A. DeSmidt, et al., 2015 A missense mutation in DCDC2 causes human recessive deafness DFNB66, likely by interfering with sensory hair cell and supporting cell cilia length regulation. Hum. Mol. Genet. 24: 2482–2491. https://doi.org/10.1093/hmg/ddv009

Hearn T., G. L. Renforth, C. Spalluto, N. A. Hanley, K. Piper, et al., 2002 Mutation of ALMS1, a large gene with a tandem repeat encoding 47 amino acids, causes Alström syndrome. Nat. Genet. 31: 79–83. https://doi.org/10.1038/ng874

Hearn T., C. Spalluto, V. J. Phillips, G. L. Renforth, N. Copin, et al., 2005 Subcellular localization of ALMS1 supports involvement of centrosome and basal body dysfunction in the pathogenesis of obesity, insulin resistance, and type 2 diabetes. Diabetes 54: 1581–1587.

Hearn T., 2019 ALMS1 and Alström syndrome: a recessive form of metabolic, neurosensory and cardiac deficits. J. Mol. Med. 97: 1–17. https://doi.org/10.1007/s00109-018-1714-x

Huang P., and A. F. Schier, 2009 Dampened Hedgehog signaling but normal Wnt signaling in zebrafish without cilia. Development 136: 3089–98. https://doi.org/10.1242/dev.041343

Imtiaz A., I. A. Belyantseva, A. J. Beirl, C. Fenollar-Ferrer, R. Bashir, et al., 2018 CDC14A phosphatase is essential for hearing and male fertility in mouse and human. Hum. Mol. Genet. 27: 780–798. https://doi.org/10.1093/hmg/ddx440

Jagger D., G. Collin, J. Kelly, E. Towers, G. Nevill, et al., 2011 Alström Syndrome protein ALMS1 localizes to basal bodies of cochlear hair cells and regulates cilium-dependent planar cell polarity. Hum. Mol. Genet. 20: 466–481. https://doi.org/10.1093/hmg/ddq493

Jones C., V. C. Roper, I. Foucher, D. Qian, B. Banizs, et al., 2008 Ciliary proteins link basal body polarization to planar cell polarity regulation. Nat. Genet. 40: 69–77. https://doi.org/10.1038/ng.2007.54

Khateb S., L. Zelinger, L. Mizrahi-Meissonnier, C. Ayuso, R. K. Koenekoop, et al., 2014 A homozygous nonsense CEP250 mutation combined with a heterozygous nonsense C2orf71 mutation is associated with atypical Usher syndrome. J. Med. Genet. 51: 460–469. https://doi.org/10.1136/jmedgenet-2014-102287

Kindt K. S., G. Finch, and T. Nicolson, 2012 Kinocilia mediate mechanosensitivity in developing zebrafish hair cells. Dev. Cell 23: 329–341. https://doi.org/10.1016/j.devcel.2012.05.022

Li G., R. Vega, K. Nelms, N. Gekakis, C. Goodnow, et al., 2007 A role for Alström syndrome protein, Alms1, in kidney ciliogenesis and cellular quiescence. PLoS Genet. 3: e8. https://doi.org/10.1371/journal.pgen.0030008

Li P., Y. He, G. Cai, F. Xiao, J. Yang, et al., 2018 CCDC114 is mutated in patient with a complex phenotype combining primary ciliary dyskinesia, sensorineural deafness, and renal disease. J. Hum. Genet. 64: 39–48. https://doi.org/10.1038/s10038-018-0514-z

Lim D. J., and M. Anniko, 1985 Developmental morphology of the mouse inner ear. A scanning electron microscopic observation. Acta Otolaryngol. Suppl. 422: 1–69.

Lindsey S., C. Brewer, O. Stakhovskaya, H. J. Kim, C. Zalewski, et al., 2017 Auditory and otologic profile of Alström syndrome: Comprehensive single center data on 38 patients. Am. J. Med. Genet. Part A 173: 2210–2218. https://doi.org/10.1002/ajmg.a.38316

Malicki J., A. F. Schier, L. Solnica-Krezel, D. L. Stemple, S. C. F. Neuhauss, et al., 1996 Mutations affecting development of the zebrafish ear. Development 123: 275–283. https://doi.org/10.5167/uzh-222

Marshall J. D., M. D. Ludman, S. E. Shea, S. R. Salisbury, S. M. Willi, et al., 1997 Genealogy, natural history, and phenotype of Alström Syndrome in a large Acadian kindred and three additional families. Am. J. Med. Genet. 73: 150–161.

Marshall J. D., R. T. Bronson, G. B. Collin, A. D. Nordstrom, P. Maffei, et al., 2005 New Alström syndrome phenotypes based on the evaluation of 182 cases. Arch. Intern. Med. 165: 675–683.

May-Simera H. L., A. Ross, S. Rix, A. Forge, P. L. Beales, et al., 2009 Patterns of expression of Bardet-Biedl syndrome proteins in the mammalian cochlea suggest noncentrosomal functions. J. Comp. Neurol. 514: 174–88. https://doi.org/10.1002/cne.22001

Meyers J. R., R. B. MacDonald, A. Duggan, D. Lenzi, D. G. Standaert, et al., 2003 Lighting up the senses: FM1-43 loading of sensory cells through nonselective ion channels. J. Neurosci. 23: 4054–4065.

Michaud J. L., E. Heon, F. Guilbert, J. Weill, B. Puech, et al., 1996 Natural history of Alstrom syndrome in early childhood: Onset with dilated cardiomyopathy. J. Pediatr. 128: 225–229. https://doi.org/10.1016/S0022-3476(96)70394-3

Nadol J. B., J. D. Marshall, and R. T. Bronson, 2015 Histopathology of the human inner ear in Alström’s syndrome. Audiol. Neurotol. 20: 267–272. https://doi.org/10.1159/000381935

Namburi P., R. Ratnapriya, S. Khateb, C. H. Lazar, Y. Kinarty, et al., 2016 Bi-allelic Truncating Mutations in CEP78, Encoding Centrosomal Protein 78, Cause Cone-Rod Degeneration with Sensorineural Hearing Loss. Am. J. Hum. Genet. 99: 777–784. https://doi.org/10.1016/j.ajhg.2016.07.010

Nesmith J. E., T. L. Hostelley, C. C. Leitch, M. S. Matern, S. Sethna, et al., 2019 Genomic knockout of alms1 in zebrafish recapitulates Alström syndrome and provides insight into metabolic phenotypes. Hum. Mol. Genet. 28: 2212–2223. https://doi.org/10.1093/hmg/ddz053

Nicolson T., A. Rüsch, R. W. Friedrich, M. Granato, J. P. Ruppersberg, et al., 1998 Genetic Analysis of Vertebrate Sensory Hair Cell Mechanosensation: the Zebrafish Circler Mutants. Neuron 20: 271–283. https://doi.org/10.1016/S0896-6273(00)80455-9

Nicolson T., 2015 Ribbon synapses in zebrafish hair cells. Hear. Res. 330: 170–177.

Nicolson T., 2017 The genetics of hair-cell function in zebrafish. J. Neurogenet. 31: 102–112. https://doi.org/10.1080/01677063.2017.1342246

Nikopoulos K., P. Farinelli, B. Giangreco, C. Tsika, B. Royer-Bertrand, et al., 2016 Mutations in CEP78 Cause Cone-Rod Dystrophy and Hearing Loss Associated with Primary-Cilia Defects. Am. J. Hum. Genet. 99: 770–776. https://doi.org/10.1016/j.ajhg.2016.07.009

Ogun O., and M. Zallocchi, 2014 Clarin-1 acts as a modulator of mechanotransduction activity and presynaptic ribbon assembly. J. Cell Biol. 207: 375–91. https://doi.org/10.1083/jcb.201404016

Owens K. N., F. Santos, B. Roberts, T. Linbo, A. B. Coffin, et al., 2008 Identification of genetic and chemical modulators of zebrafish mechanosensory hair cell death. PLoS Genet. 4: e1000020. https://doi.org/10.1371/journal.pgen.1000020

Raible D. W., and G. J. Kruse, 2000 Organization of the lateral line system in embryonic zebrafish. J. Comp. Neurol. 421: 189–198.

Reiners J., K. Nagel-Wolfrum, K. Jürgens, T. Märker, and U. Wolfrum, 2006 Molecular basis of human Usher syndrome: deciphering the meshes of the Usher protein network provides insights into the pathomechanisms of the Usher disease. Exp. Eye Res. 83: 97–119. https://doi.org/10.1016/j.exer.2005.11.010

Riazuddin S., I. A. Belyantseva, A. P. J. Giese, K. Lee, A. A. Indzhykulian, et al., 2012 Alterations of the CIB2 calcium-and integrin-binding protein cause Usher syndrome type 1J and nonsyndromic deafness DFNB48. Nat. Genet. 44: 1265–1271. https://doi.org/10.1038/ng.2426

Ross A. J., H. May-Simera, E. R. Eichers, M. Kai, J. Hill, et al., 2005 Disruption of Bardet-Biedl syndrome ciliary proteins perturbs planar cell polarity in vertebrates. Nat. Genet. 37: 1135–1140. https://doi.org/10.1038/ng1644

Rossi A., Z. Kontarakis, C. Gerri, H. Nolte, S. Hölper, et al., 2015 Genetic compensation induced by deleterious mutations but not gene knockdowns. Nature 524: 230–233. https://doi.org/10.1038/nature14580

Scheffer D. I., J. Shen, D. P. Corey, and Z. Y. Chen, 2015 Gene expression by mouse inner ear hair cells during development. J. Neurosci. 35: 6366–6380. https://doi.org/10.1523/JNEUROSCI.5126-14.2015

Sedykh I., J. J. TeSlaa, R. L. Tatarsky, A. N. Keller, K. A. Toops, et al., 2016 Novel roles for the radial spoke head protein 9 in neural and neurosensory cilia. Sci. Rep. 6: 34437. https://doi.org/10.1038/srep34437

Seiler C., and T. Nicolson, 1999 Defective calmodulin-dependent rapid apical endocytosis in zebrafish sensory hair cell mutants. J. Neurobiol. 41: 424–434.

Seiler C., K. C. Finger-Baier, O. Rinner, Y. V. Makhankov, H. Schwarz, et al., 2005 Duplicated genes with split functions: Independent roles of protocadherin 15 orthologues in zebrafish hearing and vision. Development 132: 615–623. https://doi.org/10.1242/dev.01591

Sipe C. W., and X. Lu, 2011 Kif3a regulates planar polarization of auditory hair cells through both ciliary and non-ciliary mechanisms. Development 138: 3441–3449. https://doi.org/10.1242/dev.065961

Söllner C., G.-J. Rauch, J. Siemens, R. Geisler, S. C. Schuster, et al., 2004 Mutations in cadherin 23 affect tip links in zebrafish sensory hair cells. Nature 428: 955–959. https://doi.org/10.1038/nature02484

Stawicki T. M., K. N. Owens, T. Linbo, K. E. Reinhart, E. W. Rubel, et al., 2014 The zebrafish merovingian mutant reveals a role for pH regulation in hair cell toxicity and function. Dis. Model. Mech. 7: 847–856. https://doi.org/10.1242/dmm.016576

Stawicki T. M., L. Hernandez, R. Esterberg, T. Linbo, K. N. Owens, et al., 2016 Cilia-associated genes play differing roles in aminoglycoside-induced hair cell death in zebrafish. G3 Genes, Genomes, Genet. 6: 2225–2235. https://doi.org/10.1534/g3.116.030080

Stawicki T. M., T. Linbo, L. Hernandez, L. Parkinson, D. Bellefeuille, et al., 2018 The role of retrograde intraflagellar transport genes in aminoglycoside-induced hair cell death. Biol. Open. https://doi.org/10.1242/bio.038745

Tsujikawa M., and J. Malicki, 2004 Intraflagellar Transport Genes Are Essential for Differentiation and Survival of Vertebrate Sensory Neurons. Neuron 42: 703–716. https://doi.org/10.1016/S0896-6273(04)00268-5

Van Camp G, Smith RJH, Hereditary Hearing Loss Homepage. https://hereditaryhearingloss.org (Accessed November 2, 2020)

Vona B., J. Doll, M. A. H. Hofrichter, T. Haaf, and G. K. Varshney, 2020 (online publication ahead of print) Small fish, big prospects: using zebrafish to unravel the mechanisms of hereditary hearing loss. Hear. Res. https://doi.org/10.1016/j.heares.2020.107906

Whitfield T. T., M. Granato, F. J. M. Van Eeden, U. Schach, M. Brand, et al., 1996 Mutations affecting development of the zebrafish inner ear and lateral line. Development 123: 241–254.

